# Compounding the disturbance: Family forest owner reactions to invasive forest insects

**DOI:** 10.1101/590331

**Authors:** Marla Markowski-Lindsay, Mark E. Borsuk, Brett J. Butler, Matthew J. Duveneck, Jonathan Holt, David B. Kittredge, Danelle Laflower, Meghan Graham MacLean, David Orwig, Jonathan R. Thompson

**Affiliations:** Family Forest Research Center, Department of Environmental Conservation, University of Massachusetts – Amherst, Amherst, MA; Civil and Environmental Engineering, Duke University, Durham, NC 1Harvard Forest, Harvard University, Petersham, MA; Family Forest Research Center, USDA Forest Service Northern Research Station, Amherst, MA; New England Conservatory, Boston MA; Harvard Forest, Harvard University, Petersham, MA

## Abstract

Invasive forest insect and pathogens (FIP) are having significant, direct, adverse impacts. Interactions between FIPs and forest owners have the potential to create ecosystem impacts that compound direct impacts. We assessed family forest owners’ responses to numerous contingent behavior, FIP-outbreak scenarios in the northeastern U.S. based on FIP outbreak attributes. Sixty-two percent of scenario responses (*n*=2,752) reflected a harvest intent as a result of FIPs; 84% of respondents (*n=*688) would consider harvesting in at least one of the four hypothetical scenarios presented to them. Harvest intention increased with greater FIP-related tree mortality and decreased with delayed total tree mortality. Owners with larger holdings, who had previously harvested forest products, and live on their forestland had greater intentions to harvest in response to FIPs. Results suggest that FIPs could transform the regional harvest regime with socio-ecological impacts that are distinct from those caused by FIPs or harvesting alone.

## INTRODUCTION

Forest insects and pathogens (FIPs) have significant impacts on forests worldwide. In North America, the annual area affected by FIPs exceeds that of all wildfires (Hicke et al., 2012) and in the northeastern U.S., FIPs damaged over 8 million ha during the past 17 years (Kosiba et al., 2018). Climate change and global trade are increasing the spread and severity of FIPs (Ayres & Lombardero, 2000) and the number of invasive wood-boring insects in North America is projected to increase three-to-four-fold by 2050 (Leung, Springborn, Turner, & Brockerhoff, 2014). Thus, there is urgent need to understand the full impacts of FIPs. Many of the direct impacts have been well-studied; they include: selective mortality of tree species thereby altering forest structure and composition; disruption of carbon, water, and nutrient cycles; and reduction in ecosystem service provision including timber production, carbon storage, and habitat (See reviews by: Lovett et al., 2016; Peltzer, Allen, Lovett, Whitehead, & Wardle, 2010).

Less well-studied are the human-mediated indirect impacts of FIPs. Specifically, the presence of FIPs often instigates salvage logging. Indeed, even the threat of FIPs can trigger pre-emptive logging. The effects of logging may generate more profound ecosystem disruption and impacts on biodiversity than the FIP itself (Foster & Orwig, 2006; Thorn et al., 2018). Salvage logging is a common management response to forest disturbance with distinct ecological consequences that are increasingly recognized (Lindenmayer, Burton, & Franklin, 2012). Ecologically, the salvage response to FIPs can be considered a compounding disturbance that intensifies many aspects of the disturbance and often broadens the number of affected tree species (e.g., through harvest “by-catch” of more merchantable species to defray harvest costs). A full accounting of the direct and indirect impacts of FIPs mediated by the human response requires a coupled human and natural systems perspective and, specifically, a much better understanding of forest owner response.

Although many stakeholders influence the management response to FIPs, private forest owners are an essential group. Private forest owners control the plurality (58%) of forestland in the U.S.; approximately 36% is held by an estimated 11-million families, individuals, trusts, estates, and family partnerships, collectively referred to as family forest owners (FFOs) (Butler et al., 2016a). FFOs make independent forest management decisions as they see fit. In FFO-dominated landscapes, policy-makers and conservationists are challenged by the “tyranny of small decisions,” wherein the aggregation of many small independent decisions determine the regional-scale ecological outcomes without any explicit consideration of the broader context (Odum, 1982). Despite the widely acknowledged need for empirical studies of human behavior and decision-making regarding land management and conservation (Cowling, 2014; Field, Dayer, & Elphick, 2017), few studies have investigated the management response to FIPs.

Studies of forest owner response to FIPs have focused on the southern pine beetle (SPB), which has infested much of the southeastern U.S. These studies have characterized owners’ willing to consider pre-emptive or salvage control measures and focused on approaches to motivate forest management and promote forest health (Mayfield III, Nowak, & Moses, 2006; Molnar, Schelhas, & Holeski, 2007). Here, harvesting was more likely among those having experience with forest management professionals, greater losses associated with SPB (Molnar et al., 2007) and larger landholdings (Mayfield III et al., 2006). SPB is an important species in the South (Schleeweis et al., 2013), but the number of different FIPs with unique, damaging attributes is growing worldwide (Liebhold et al., 2017). The SPB-focus of these studies precludes an understanding of other FIP attributes that are important to owners when considering management.

Here we investigate the FFO response to FIPs in the northeastern U.S., an ideal land system to study the human dimensions of FIP infestations. New England is among the most forested and populous regions of North America. Approximately 84% of forests are privately owned and 41% are owned by FFOs (Butler et al., 2016b). While wood products are not a dominant sector of the regional economy, partial forest harvesting is the dominant ecological disturbance, with attributes of the harvest regime (i.e., frequency and intensity of cutting) jointly controlled by social and biophysical factors (Thompson, Canham, Morreale, Kittredge, & Butler, 2017). FFOs in the region tend to own their land primarily for privacy and aesthetic reasons, but many harvest trees commercially (Butler et al., 2016a). Forest land management occurs infrequently and is triggered by poorly-understood, exogenous events (Kittredge, 2004; Markowski-Lindsay, Catanzaro, Milman, & Kittredge, 2016). Most owners do not have written forest management plans and have not received professional advice (Butler et al., 2016b). The Northeast also has the highest diversity of non-native FIPs in the U.S. (Figure 1). The leading FIP species of concern include the hemlock woolly adelgid (*Adelges tsugae*), emerald ash borer (*Agrilus planipennis*), and gypsy moth (*Lymantria dispar*), which all have different host tree species with distinct ecological functions and services.

**Figure 1.**
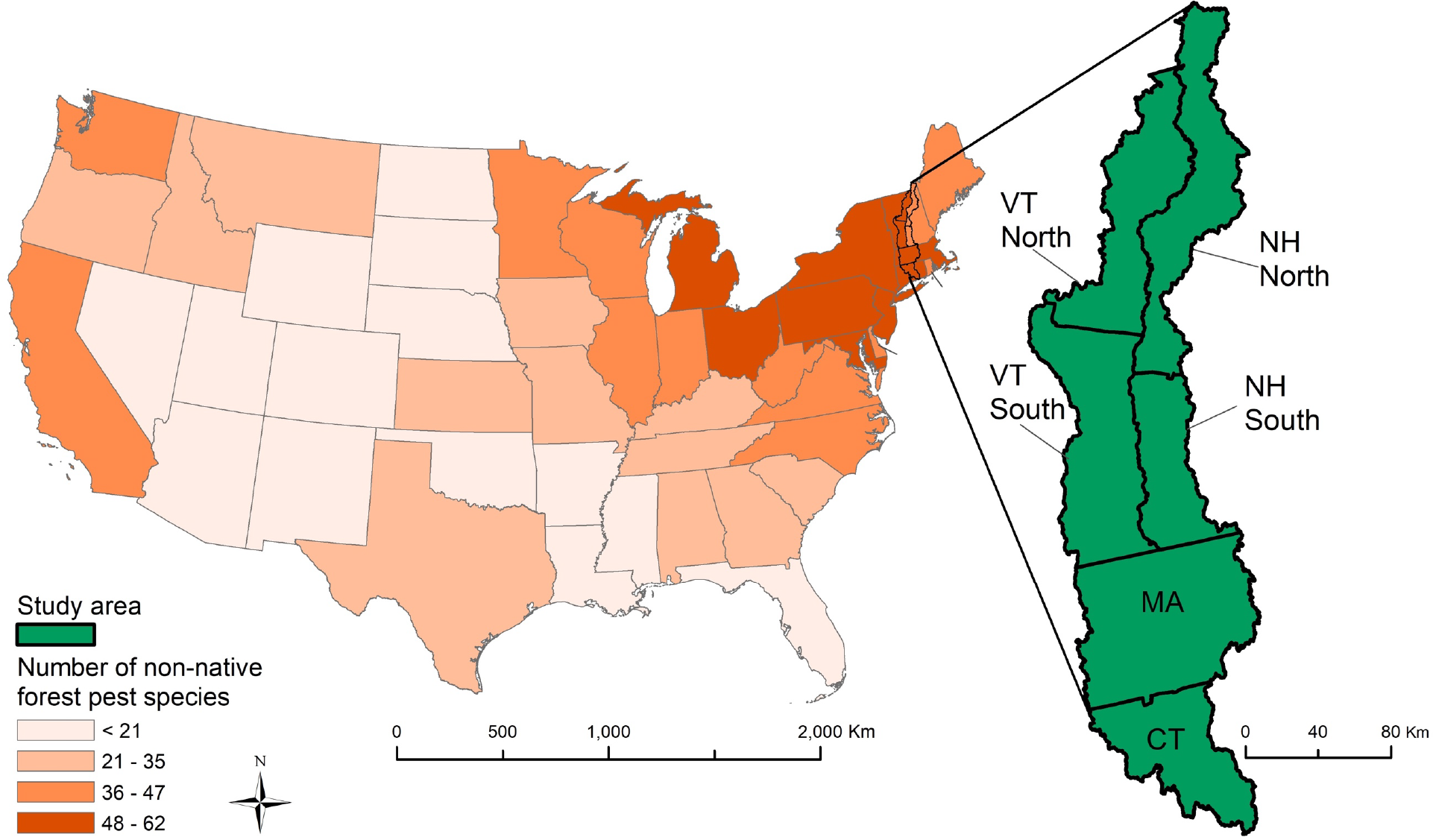
The Connecticut River watershed study area and distribution of non-native forest insect pest species in the conterminous U.S. in 2015 (Adapted from: Cary Institute of Ecosystem Studies, 2019).

We hypothesize that a widespread FIP outbreak could serve to instigate harvest decisions resulting in regional-scale, possibly synchronized, socio-ecological consequences that exceed those of the FIP alone. To begin to test our hypothesis, we surveyed FFOs to understand the circumstances under which FIP infestation would induce them to harvest. Our results suggest that future outbreaks of FIPs in the Northeast could alter the regional harvest regime, especially when FIPs cause high levels of tree mortality.

## METHODS AND ANALYSIS

### Sample selection

We surveyed FFOs owning ≥ 4 ha within the 2.9 million ha Connecticut River watershed stratifying across six state/sub-state regions (Connecticut, Massachusetts, north and south New Hampshire, and north and south Vermont) (Figure 1) and by parcel sizes of 4 – 19 ha and ≥ 20 ha, to ensure larger parcels were represented in the sample. We randomly selected FFOs within each stratum from property tax records.

### Survey design and administration

To assess likely FFO response to FIPs, we constructed a series of contingent behavior questions (Englin & Cameron, 1996) based on four key FIP attributes at varying levels: (i) arrival time, how virulent or damaging the FIP is, including (ii) tree mortality percentage, (iii) time until 100% tree mortality, and (iv) value of timber loss. We selected these attributes to reflect characteristics of the main FIPs in the northeast (Lovett, Canham, Arthur, Weathers, & Fitzhugh, 2006). Each respondent was presented with four infestation scenarios based on these generic FIP attributes and asked whether they would harvest trees targeted by the FIP and how certain they were of their response (See Text S1, Supporting Information). The survey also asked respondents to provide information on ownership characteristics.

In 2017 we sent 2,000 mail surveys to approximately 333 FFOs per strata, following methods described by Dillman, Smyth, & Christian (2014). We obtained a 37% cooperation rate and based on follow-up telephone calls, detected no evidence of nonresponse biases. We imputed item nonresponse values using a random forests approach (*sensu* Stekhoven & Buhlmann, 2012).

### Decision to harvest trees targeted by FIPs

A multilevel regression model (MLM) (Gelman & Hill, 2007) was used to account for the multiple scenario responses from each respondent. We established two separate MLMs of FIP-induced harvesting intention that incorporated respondent uncertainty: a base model relating the four attributes of the hypothetical FIP to harvest intention, and an expanded model supplementing the base model with owner-specific characteristics. We incorporated weights into the models to account for the stratified sample design. The weights, *ω*_*s*_, were a function of the population size, *N*_*s*_, and number of respondents, *n*_*r,s*_, in each stratum, *s*, (i.e., *ω*_*s*_ = *N*_*s*_ / *n*_*r,s*_).

The base model described the probability of harvesting trees affected by the hypothetical FIP depending on the four FIP attributes varied in each contingent behavior question. We hypothesized that harvest intention would vary with FIP attributes. FIP attributes were coded as discrete categories, so that the model results would indicate the differences in harvest intent across the levels of the FIP attributes (Table S2).

The expanded model included owner characteristics, and we hypothesized that, in addition to FIP attributes, harvest intentions would vary with demographics (i.e., age, gender, education, income, absenteeism, land tenure, parcel size), harvesting familiarity (i.e., history of past harvest and of receiving professional advice), and ownership objectives (i.e., owning for timber production, investment, consumption, protection, and recreation (Table S1, Supporting Information). These characteristics, taken from the survey, correspond to those in FFO harvesting literature (see review by Silver, Leahy, Weiskittel, Noblet, & Kittredge, 2015). To characterize passive owners (*sensu* Silver et al., 2015), we also included whether the owner was likely to transfer their land in the near future. Finally, to examine whether the presence of a FIP in a respondent’s town would induce harvest, we included whether two common FIPs had been detected in the respondent’s town. We tested for multicollinearity among potential explanatory variables using Variance Inflation Factor (VIF) diagnostics; VIF tolerance levels below 0.4 are associated with high multicollinearity (Allison, 1999). The lowest level for variables in this analysis was 0.5.

Both models incorporated respondent uncertainty, which we measured by following the scenario questions with one that asked respondents to rate how certain they were of their answer, on a five-point Likert scale. We used the symmetrical uncertainty model as a guide by mapping responses to a numerical certainty scale (Loomis & Ekstrand, 1998). The numerical scaling projects a range of certainty onto the binary responses, with the lower end of the scale reflecting more certain negative responses, the middle scale the most uncertain responses, and the upper scale increasing certainty in positive responses (See Table 1).

**Table 1.**
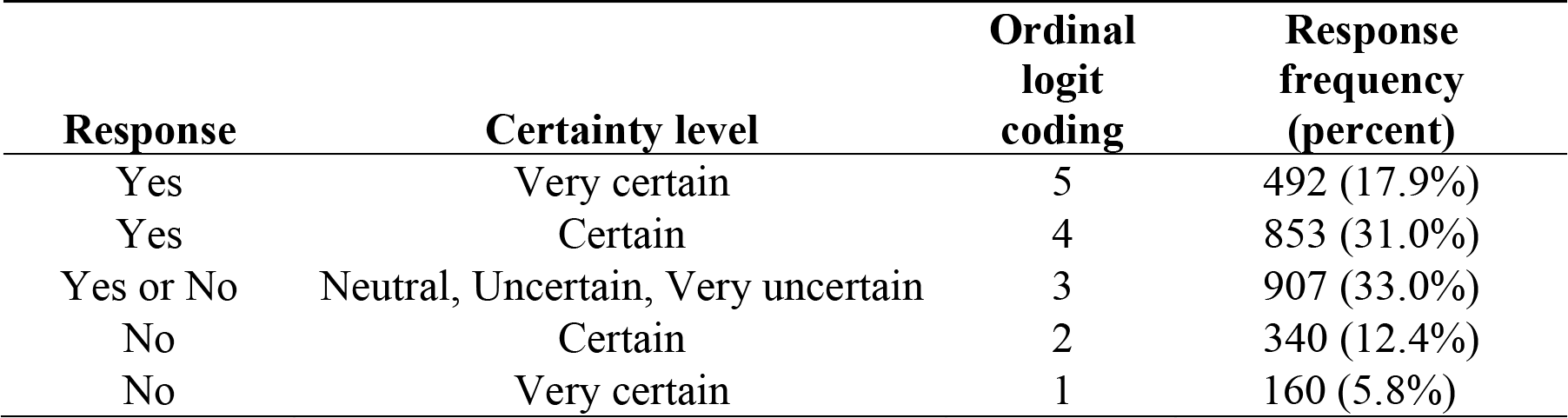
Scenario response coding for uncertainty (*n* = 2,752)

For both models, we fit the data using ordinal logistic MLM (Greene, 2011) on the harvest intention coded to account for uncertainty. We used the Stata15 *meologit* package using the *svyset* option to specify the survey design sampling units (i.e., respondents) and weights.

## RESULTS

Of the 2,000 surveys, 688 respondents provided usable surveys. The models, accounting for correlation among the four scenario responses for each respondent, thus analyzed 2,752 scenario responses. Nearly 50% of scenario responses were either certain or very certain about the intent to harvest in response to FIPs while nearly 20% of scenario responses were certain or very certain about the intent not to harvest (Table 1).

While we model responses, it is informative to consider how *respondents* replied to scenarios; cut intention for 63% of respondents varied with FIP attributes or reflected respondent uncertainty (Table 2). In fact, 84% of respondents would consider harvesting for at least one of the four scenarios presented.

**Table 2.** Cut intention by respondent (*n* = 688)

| **Response** | **Response frequency**<br>**(percent)** |
| --- | --- |
| Respondent would cut under every scenario and is *Very certain* or *Certain* about each of these responses | 196 (28.5%) |
| Respondent cut intention and/or certainty varies across scenarios | 435 (63.2%) |
| Respondent would not cut under every scenario and is *Very certain* or *Certain* about each of these responses | 57 (8.3%) |

The majority of respondents were male, lived on their land, had a college degree, had experience cutting timber for commercial purposes, owned their land for the objective of protection, less often owned their land for investment or timber objectives, and were infrequently apt to sell or give away their land in the next 5 years (Figure 2). Roughly half of respondents owned their land for consumptive or recreation objectives, and have received professional advice on the care, management or protection of their land. While the majority earned less than $100,000 per year, nearly 44% earned more than this. Nearly 45% lived in a town where one of the two leading FIP species of concern had already been detected. Respondents, on average, own 44 ha of wooded land, have owned their land for nearly 25 years, and are nearly 65 years of age. (See Figure 2)

**Figure 2.**
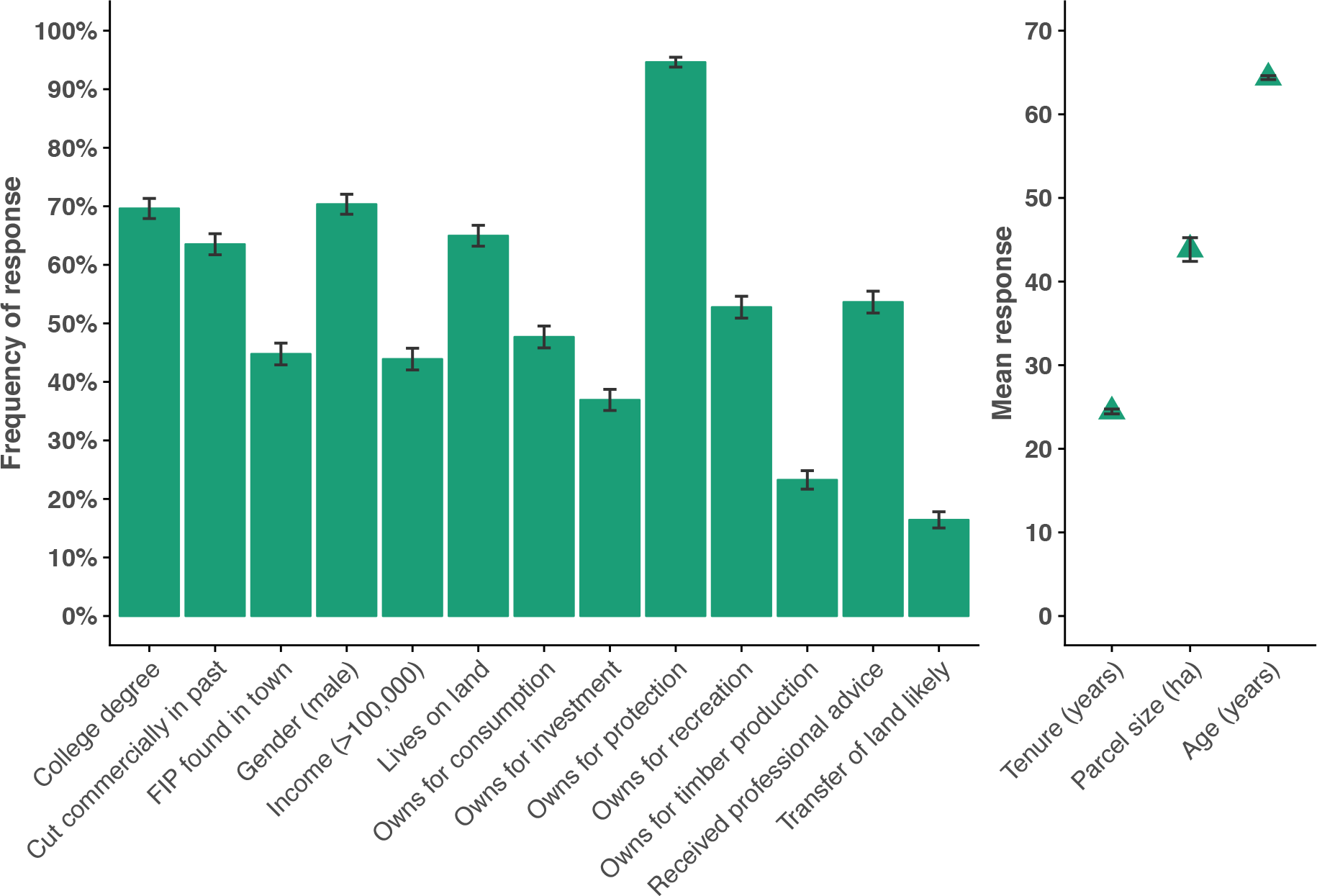
Family forest owner characteristics frequency/means and 95% confidence intervals. The left panel shows discrete owner characteristic frequencies while the right shows continuous owner characteristic means.

### Models

FIP attribute coefficients and significance levels in the base and expanded models were nearly identical. Individually, both had significant F-statistics and the adjusted Wald test indicates significance (Table S2, Supporting Information).

The odds ratios indicate how the FIP attribute levels influence the harvesting intent likelihood. Respondents are more likely to intend to harvest when FIPs kill a greater proportion of their trees (i.e., greater mortality effect; Figure 3). A mortality effect of 50% increases the odds of harvest intention by a factor of 2.8 over a mortality effect of 10%, whereas a 90% mortality effect increases the odds of harvest intention by 4.2 over a 10% mortality effect. Harvest intention is also positively related to greater tree value loss. While respondent harvest intention does not differ significantly between a tree loss value of 50% and 10%, the odds of harvest increase by 1.6 with 90% value losses over 10% losses. Respondents are less likely to intend to cut the longer it takes for the FIP to kill the trees: increasing the time it takes for a FIP to kill all trees from 5 to 15 years decreases the odds of harvest intention by 0.73. Arrival time had no effect on intention to cut in this model. (See Figure 3; Table S2)

**Figure 3.**
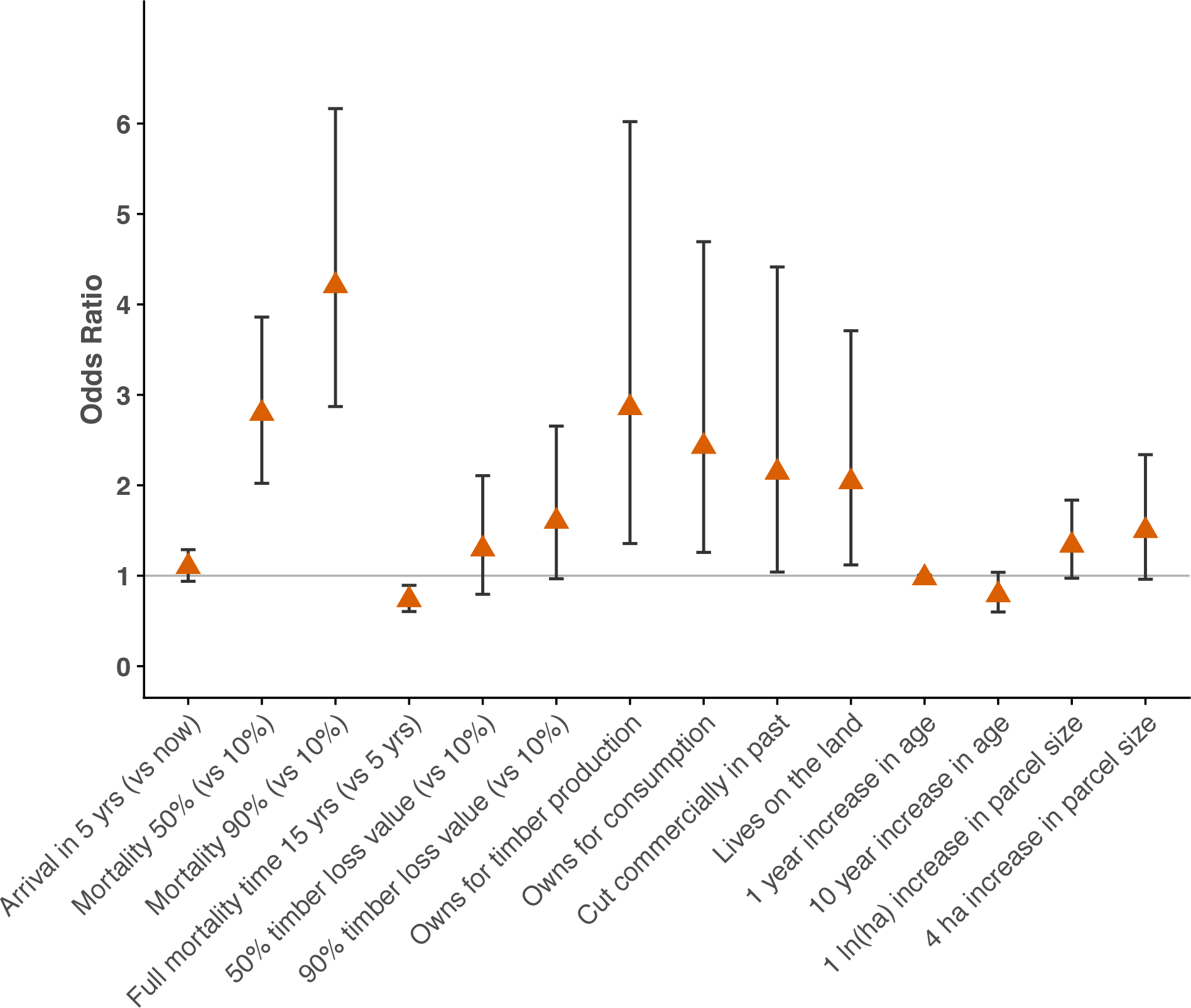
Odds ratio results (with 95% confidence intervals) for FIP attributes (coded as discrete categories) and significant family forest owner characteristics (p ≤ 0.10).

Owner characteristics with the highest likelihood of harvest intention include: owning for timber and consumption, having past commercial harvests, and living on the land that they manage. Having these characteristics (versus not) increases the odds of harvest intention by more than a factor of 2. Two continuous factors influence FIP-induced harvest intention at lower significance levels: age and parcel size. Older respondents were less likely to cut; every 10 additional years in age decreases the odds of harvest intention to 0.8. Those owning more land are more likely to cut. Every 4 ha increase in land ownership raises the odds of harvest intentions by 1.5 (Figure 3). The direction of these results are consistent with prior FFO harvesting studies (e.g., Aguilar, Cai, & Butler, 2017; Silver et al., 2015).

## DISCUSSION

The majority of FFOs we surveyed indicated that their intent to harvest varied with FIP attributes. Similar to a southern U.S. study (Molnar et al., 2007), we found FIP mortality effects influenced harvest intention, and this influence was greatest of all attributes. The higher the percent of trees killed by the FIP, the greater the likelihood of harvest. While the timing of FIP arrival did not matter to respondents in their intent to harvest, a delay in the time for all trees to die decreased harvest intention. FIP arrival timing did not impact intent to harvest. Similarly, the value of timber lost did not greatly impact harvest intention confirming prior findings of how timber value does not motivate harvesting in the region (Kittredge & Thompson, 2015).

If these harvest intentions are borne out, the regional-scale consequences of FIP infestations—in terms of effects on commodity production, ecosystem services, and biodiversity protection— would be altogether distinct from FIPs infestations alone. The effects of salvage logging are ecologically distinct (Thorn et al., 2018) and FIP-induced salvage harvests have particularly compounding impacts because they often remove trees and tree species that were not affected by the initial disturbance. Also, FIP infestations are a unique harvesting trigger. Forest management decisions are motivated by various reasons and are largely uncoordinated (Kittredge, 2004; Markowski-Lindsay et al., 2016). However, FIP infestations potentially offer region-wide synchronizing event among some FFOs, particularly when mortality impacts are high. Certainly, a coordinated, synchronized harvesting event may be limited by poor wood markets or expense (Mayfield III et al., 2006), biophysical and social availability of wood from family forests (Butler, Ma, Kittredge, & Catanzaro, 2010), or other factors. Nonetheless, direct FIP ecosystem impacts may be exacerbated by such synchronicity, be they local or region-wide, creating a coupled natural- and human-disturbance system with even greater ecosystem repercussions than each one individually. FIP disturbances are growing with time (Ayres & Lombardero, 2000) and as the spread and – especially – severity increase with climate change, it is critical to understand the human response at national and global scales.

While it remains to be tested, our results suggest that FIP-induced harvest intention may be transferrable to other regions, especially those dominated by FFOs with similar characteristics. Here, FFOs with physically close ties to their land (i.e., experience with and goals of timber production, objectives of woodland resource consumption, and resident owners) had the greatest likelihood of harvesting, consistent with extant studies (e.g., Aguilar et al., 2017; Molnar et al., 2007; Silver et al., 2015). Across the U.S., 22% of ownerships own their land for timber, 29% have cut experience with commercial cutting, 44% for own for consumption purposes, and 63% of the ownerships were non-absentee (Butler et al., 2016b). Focusing specifically on the owners who own their land for timber (22% of all FFOs in the U.S.), our results suggest that FIP-induced harvest would be more likely on these roughly 94 million acres of forest, reflecting 37% of all FFO-owned land, then land owned by FFOs not having this ownership objectives. Focusing on resident FFOs (63% of all FFOs in the U.S.), our results suggest that FIP-induced harvest would be more likely on these roughly 148 million acres of forest, reflecting 56% of all FFO-owned land, then land owned by FFOs who do not live on their land.

Our next steps in assessing the complex feedbacks within this natural- and human-system will be to use these results in simulations designed to represent FFO behaviors as they are confronted with the presence or threat of infestation. These simulations will estimate regional-scale changes in forest structure, carbon and species composition as they are affected by FIP dynamics, climate change, and the land-use regimes articulated via FFO behavior models. The coupling between the behavioral model representing the human system and the forest simulation representing the natural system should be dynamic to capture the specific patterns that emerge from the complex feedbacks between the two. Additional research is also needed to understand how FIPs affect management and harvesting practices (*sensu* Molnar et al., 2007). Incorporating field and social science data into simulations can be used to better quantify the long-term and broad-scale impacts of FIPs on future forest conditions and to identify strategies that best conserve and sustain multiple ecosystem services and conservation values. Additional complexities to be pursued include better understanding of the marketplace system, including the impact flooded wood markets may have on decisions or the effect quarantines may have on timber transportation and the capability of loggers to respond to FFO harvest decisions.

## Acknowledgements and Data

This material is based on work supported by the National Science Foundation under Grant No. DEB-1617075. This research is partially supported by the Harvard Forest Long Term Ecological Research Program Grant No. NSF-DEB 12-37491). The survey questionnaire and procedures were approved by the lead author’s Institutional Review Board (IRB) in accordance with the Human Research Protection Program. [The human subjects data used in this manuscript will be forthcoming in a limited format to comply with IRB requirements before publication.]

## SUPPORTING INFORMATION

**Table S1.** Model variables, definitions and coding

| Variable | Definition |
| --- | --- |
| <u>FIP attributes</u> |  |
| Arrival time | Now (0), in five years (5) |
| Mortality percentage | Percent of trees killed: 10%, 50%, 90% |
| Time until full mortality is reached | Years to mortality: 5, 15 |
| Value of timber loss | Percent of economic value loss: 10%, 50%, 90% |
| <u>Expanded model characteristics</u> |  |
| Age | Years (continuous) |
| College degree | Has college degree: Yes (1), No (0) |
| Cut commercially in past | Yes (1), No/Don't know (0) |
| FIP found in town | Yes (1), No (0) |
| Gender (male) | Male (1), Female (0) |
| Income | Annual income \$100,000 or more: Yes (1), No (0) |
| Lives on land | Lives within 1 mile of land: Yes (1), No (0) |
| Owens for consumption<sup>a</sup> | Very important/important (1), Otherwise (0) |
| Owens for investment<sup>a</sup> | Very important/important (1), Otherwise (0) |
| Owens for protection<sup>a</sup> | Very important/important (1), Otherwise (0) |
| Owens for recreation<sup>a</sup> | Very important/important (1), Otherwise (0) |
| Owens for timber production<sup>a</sup> | Very important/important (1), Otherwise (0) |
| Parcel size | Size of woodland owned (hectares, logged) |
| Received professional advice<sup>b</sup> | Yes (1), No/Don't know (0) |
| Tenure | Number of years owning land (continuous) |
| Transfer of land likely in 5 years<sup>c</sup> | Extremely likely/likely (1), Otherwise (0) |

<sup>a</sup>The survey asked respondents how important various reasons are for owning their land on a 5-point Likert scale of *Very important* to *Not important*. A respondent was coded as having the objective if they responded *Very important* or *Important* to the Likert scale for that question.
- Owens for timber production: Responded *Very important* or *Important* to “For timber products, such as logs or pulpwood” - Owens for investment: Responded *Very important* or *Important* to “For land investment” - Owens for consumption: Responded *Very important* or *Important* to any of the following: “For firewood”, “For nontimber forest products”, or “For hunting” - Owens for protection: Responded *Very important* or *Important* to any of the following: “To enjoy beauty or scenery”, “To protect nature or biological diversity”, “To protect or improve wildlife habitat”, or “For privacy” - Owens for recreation: Responded *Very important* or *Important* to “For recreation, other than hunting”
<sup>b</sup>The survey asked: “Have you ever gotten advice about the care, management or protection of your woodland from a consulting forester or another professional?”
<sup>c</sup>The survey asked respondents how likely it is they would sell or give away any of their woodland in the next 5 years on a 5-point Likert scale of *Extremely likely* to *Extremely unlikely*. If they answered *Extremely likely* or *Likely* they were coded as being likely.

**Table S2.** FIP attribute model comparison results for base and expanded models.

| Independent variable | Base model<sup>a</sup> |  | Expanded model<sup>a</sup> |  |
| --- | --- | --- | --- | --- |
|  | Coefficient | Odds Ratio | Coefficient | Odds ratio |
| Arrival time 5 yrs (vs. now) | 0.10 | 1.10 | 0.09 | 1.10 |
| Mortality 50% (vs. 10%) | 1.03*** | 2.80*** | 1.03*** | 2.79*** |
| Mortality 90% (vs. 10%) | 1.44*** | 4.23*** | 1.44*** | 4.21*** |
| Time to 100% tree mortality 15 yrs (vs. 5) | -0.31** | 0.73** | -0.31** | 0.73** |
| Value of timber loss 50% (vs. 10%) | 0.26 | 1.29 | 0.26 | 1.29 |
| Value of timber loss 90% (vs. 10%) | 0.47^ | 1.60^ | 0.47^ | 1.60^ |
| Age |  |  | -0.02^ | 0.98^ |
| College degree |  |  | -0.10 | 0.90 |
| Cut commercially in past |  |  | 0.76* | 2.14* |
| FIP found in town |  |  | -0.23 | 0.79 |
| Gender (male) |  |  | 0.01 | 1.01 |
| Income |  |  | 0.33 | 1.39 |
| Lives on land |  |  | 0.71* | 2.04* |
| Owens for consumption |  |  | 0.89** | 2.43** |
| Owens for investment |  |  | 0.48 | 1.61 |
| Owens for protection |  |  | 0.12 | 1.13 |
| Owens for recreation |  |  | 0.08 | 1.08 |
| Owens for timber production |  |  | 1.05** | 2.86** |
| Parcel size (ln(hectares)) |  |  | 0.29^ | 1.34^ |
| Received professional advice |  |  | 0.47 | 1.60 |
| Tenure |  |  | 0.004 | 1.00 |
| Transfer of land likely |  |  | 0.30 | 1.35 |
| Cutoff 1<sup>b</sup> | -4.47 |  | -2.97 |  |
| Cutoff 2<sup>b</sup> | -1.84 |  | -0.34 |  |
| Cutoff 3<sup>b</sup> | 1.19 |  | 2.69 |  |
| Cutoff 4<sup>b</sup> | 4.60 |  | 6.11 |  |
| Var (constant) | 10.37 |  | 8.77 |  |

<sup>a</sup>p-value: $\leq 0.1\% = ***$ , $\leq 1\% = **$ , $\leq 5\% = *$ , $\leq 10\% = ^$
In both models, F-statistics are significant at $p < 0.00001$ (Base: $F(6, 682) = 27.16$ , Expanded: $F(22, 666) = 11.2$ ; $n=2,752$ ). Adjusted Wald test for exclusion of additional independent variables in the Expanded model is significant at: $F(16, 672) = 4.31$ , $\text{Prob} > F = 0.0000$ .
<sup>b</sup>Ordinal models are based on latent variables governing respondent choices. We assume respondents make choices that increase their utility, and there is a continuous, unobservable variable that represents opinion level or utility association with their choice. While we do not directly observe respondents' utility for each scenario, we assume a transformation function (*sensu* Klosowski, R., Stevens, T., Kittredge, D., Dennis, D., 2001. Economic incentives for coordinated management of forest land: a case study of southern New England. *Forest Policy and Economics* 2, 29–38.) that enables construction of the ordered model wherein we define unknown utility cutoffs delineating the five responses presented in Table 1 (see Greene, W. H. (2011). *Econometric Analysis* (7th ed.). Prentice Hall). If a respondent's utility from a scenario is below the first cutoff, he or she chooses response 1; if utility is between the first and second cutoffs, the response is 2, etc. The first cutoff is normalized to zero, as is standard practice, thus the model estimates the four remaining cutoffs as presented in this table.

### Text S1. Survey Design Detail

To assess likely landowner response to invasive forest insect and pathogens (FIPs), we constructed survey based on four key FIP attributes at various levels: FIP arrival time, mortality percentage, time until full mortality is reached, and value of timber loss.

- Arrival time: While some areas have FIPs and others do not, we test two FIP arrival times to get at this characteristic: current infestation and infestation in 5 years.
- Mortality percentage: While some FIPs focus on one tree species (e.g., hemlock woolly adelgid) and others are indiscriminate or have multiple hosts (e.g., gypsy moth), we capture the range of potential FIP virulence with three mortality levels: killing 10%, 50%, and 90% of the trees.
- Time until 100% mortality of host trees is reached: The time it takes for FIPs to kill trees varies; as such, we test two mortality rates: trees that would be killed within 5 or 15 years.
- Value of timber loss: Differences in host specificity and associated tree value impacted by FIPs can result in differences in economic value losses – the economic impacts of higher value ash losses likely differ from those of lower value hemlock. To test value of timber loss, we test three values: timber reduction values of 10%, 50%, and 90%.

Each respondent was presented with four FIP infestation scenarios based on the FIP attributes and asked if they (or someone they hired) would cut or remove trees targeted by the FIP for each scenario. The 36 possible scenarios were reduced to 16 using the standard fractional factorial design and then grouped into four survey versions – each version containing a distinct set of four scenarios reflecting plausible impacts based on the various FIPs in the region. For each hypothetical scenario, respondents were asked how certain they are about their answer to the question, using a 5-point Likert scale rating of certainty.

FIP attributes were coded as discrete categories, so that the model results would indicate the differences in harvest intent across the levels of the FIP attributes.

